# A weighted graph of the projections to mouse auditory cortex

**DOI:** 10.1101/228726

**Authors:** Nuno Macarico da Costa, Kevan A.C. Martin, Franziska D. Sägesser

## Abstract

The projections to individual cortical areas from extrinsic sources are a major determinant of the area’s function, but we lack comprehensive quantitative input maps even for primary sensory areas in most model species. To quantify all input sources to the mouse primary auditory cortex (Au1), we made localized injections of modified rabies virus (SADΔG-mCherry) into Au1 of five C57BL/6 mice and identified all the cortical and subcortical areas containing retrogradely labeled cells. Of all neurons projecting to Au1 from extrinsic areas, 27 % were located in the ipsilateral cortex, 14 % in the contralateral cortex, and 58 % in subcortical regions (almost exclusively ipsilateral, predominantly in the medial geniculate nucleus). Although 90 % of the labeled cells in the ipsilateral cortex were located within 1 mm of Au1, most cortical areas projected to Au1, including visual, somatosensory, motor, rhinal, cingulate and piriform cortices. The hierarchical relations of the cortical areas projecting to Au1 were determined based on the proportion of cell bodies in superficial versus deep layers. Feedback projections (from deep layers 5/6) dominated, but temporal association and auditory cortices were on the same hierarchical level, providing input from both superficial and deep layers. Au1 is embedded in a densely connected network that involves a high degree of cross-modal integration.

## Introduction

New insights into the rules of inter-areal connectivity of neocortical circuits have been given by quantitative studies of the macaque monkey where retrograde tracing techniques combined with quantitative data collection and sophisticated statistical and graph theoretic analyses have completely revolutionized our concepts of cortical wiring in a primate brain (Markov et al., 2011, 2012, 2013; Ercsey-Ravasz et al., 2013). Whereas before macaque cortex was seen as a modular organization of sensory, motor and association areas linked in a ‘small world’ network, it is now clear that the networks are not small world, but dense, and the probability of connection between areas declines exponentially with physical distance. This latter feature means that the connections between nearest neighboring areas approach all-to-all, but long-range connections form a very small fraction of the total complement of input to an area.

In the comparatively tiny brain of the mouse, such quantitative retrograde studies have not been attempted, so it remains to be discovered whether the same rules of connectivity apply. One reason for the lack of comprehensive, detailed and quantitative studies is technical: cortical areas in the macaque are large, so large pressure injections of conventional tracers can be placed with high precision. But in the mouse, even primary areas are small, e.g. area 41 (primary auditory cortex, Au1) is only 0.8 mm^2^ in area, and thus injecting sufficient amounts of conventional retrograde tracers with high accuracy is a challenge. Work on mouse cortical connectivity has revealed details of inter-areal projection maps of a number of visual areas (Wang and Burkhalter, 2007), and now a major goal is to describe the mouse ‘connectome’ by performing large-scale tracing studies (Oh et al., 2014; Zingg et al., 2014). These studies have not, however, exploited the quantitative methods used so successfully in the macaque.

In this study of the inputs to the primary auditory cortex in the mouse we adopted the methods pioneered in the macaque studies. The prime advantage of using retrograde rather than anterograde tracing methods is that counting neurons is far simpler than counting synapses. The number of retrogradely labeled neurons makes for a clearly defined measure of relative input strengths of different source areas to a target area. The cortical laminae where the projecting neurons are located provides additional information about the nature of the connection to the target area and gives hints concerning the relative hierarchical position of cortical source and target area (Rockland and Pandya, 1979; Barone et al., 2000). Thus here we have expressed the relative connection ‘weight’ of each area as the fraction of retrogradely labeled neurons (FLN, Markov et al., 2011) and further determined the fraction of labeled supragranular layer neurons (SLN, Barone et al., 2000) as a measure of ‘feedforward’ versus ‘feedback’ projections for cortical areas.

These data answer the important question of whether the principles of interareal connectivity discovered in the macaque also apply to mouse neocortex. Unlike in other mouse connectivity studies, the distribution of FLN and SLN values we found can be directly compared to the data from macaques. At a time in which the mouse is the most widely used model system in neuroscience, it is becoming more relevant to link quantitative findings on rodent brain networks and function to the extensive work in primates.

## Materials and Methods

All experiments were performed in agreement with European Communities Council Directive of November 24, 1986 (86/609/EEC), the Veterinary Office of Zurich guidelines under Licence No. 16/2010 to K.A.C. Martin and in accordance with the Basel Declaration principles. Seven young adult male C57BL/6 mice (B6CBAF1/OlaHsd (C57BL/6 x CBA), Harlan lab, Netherlands) were used for this study, five for virus injections and two for BDA injections. The adeno-associated virus AAV2/5.CMV.EGFP encoding enhanced green fluorescent protein (EGFP) was purchased from Penn Vector Core, Pennsylvania. The glycoprotein-deleted rabies virus encoding the red fluorophore mCherry [SADΔG-mCherry] (Marshel et al., 2010) was a generous gift from Botond Roska, Friedrich Miescher Institute for Biomedical Research, Basel, Switzerland (Yonehara et al., 2011).

### Surgical procedures and perfusion

After induction of anaesthesia with 4 % isoflurane in 100 % oxygen, the mouse was fixed through a lidocaine-covered mouthpiece and prepared for stereotaxic surgery. During the surgery, the isoflurane level was reduced to 2-3 % and the body temperature of the mouse was monitored and maintained at 37 °C. From the Paxinos and Franklin atlas (Paxinos and Franklin, 2001), the coordinates for the left primary auditory cortex (Au1), were chosen to be 2.9 mm posterior and 4.1 mm lateral in relation to Bregma and the injection pipette was positioned at an angle of 30 degrees from the horizontal axis. A distinct Y-shaped blood vessel immediately posterior to Au1 served as an additional reference point for the injection site. The location of injection was later confirmed in histological sections.

Mice were injected with rabies virus or a mixture of adeno-associated virus (AAV, 1/4 to 1/3 of the virus mixture) and rabies virus (3/4 to 2/3 of the virus mixture) or with biotinylated dextran amine (BDA, MW 10000, Invitrogen) through a micro-pipette of 10-20 μm inner tip diameter. AAV was used to identify the maximum extent of the injection site, and rabies as the tracer, which retrogradely labels neurons with terminals in the injection site. The exact titer of the rabies virus was unknown; but it is expected to have been up to 10^10^ infectious units per ml (Wickersham et al., 2010). A volume of 100-200 nl of the virus mixture was injected via a glass pipette by pressure. In mice 1 and 2 the injection site was centered in layer 4 and 5, but all layers (except the deepest part of layer 6) were labeled. In the other mice small volumes were injected ca. every 200 μm to label more evenly the whole depth of Au1. A second set of mice were injected iontophoretically with BDA-MW10000 dissolved 1:50 in electrophoretic solution (0.2 M KCl, 0.05 M Tris Base, pH 7.9) instead of with virus. We started injecting (2-3 μA; 5 s on, 5 s off) at ca. 800 μm depth and moved up 100 μm every 2 min, so the total injection time was about 20 min. Post-injection, the mice were kept for 7 to 10 days to allow for the spread of the viruses and the expression of the fluorescent proteins. BDA-injected mice recovered for 2 weeks.

The mice injected with rabies were then transcardially perfused under deep anesthesia (100 mg/kg sodium pentobarbital administered intraperitoneally). After 0.1 ml heparin (10 % in Ringer solution) was injected into the left ventricle of the heart to prevent blood clotting, the brain was perfused with saline (0.9 % NaCl), follwed by 4 % paraformaldehyde (PFA, 200-300 ml in total). The brain was removed and kept in 0.1 M phosphate buffer (pH 7.4) overnight or in 4 % PFA until further processing. The BDA-injected mice were perfused with 4 % PFA, 15 % picric acid and (freshly added) 0.8 % glutaraldehyde, followed by a sucrose-gradient (10 %, 20 % and 30 % sucrose) perfusion. Then the brains were sunk in 30 % sucrose.

### Preparation for microscopy

The fixed brains were embedded in 2 % agarose. For the fluorescence microscopy, the coronal sections of 60 μm thickness were cut with a Microm HM 650 V vibratome. BDA+ brains were freeze-thawed (i.e. held in a plastic beaker which was dipped in liquid nitrogen until the brain was completely white and then transferred quickly to 0.1 M phosphate buffer) and coronal sections were cut at 80 μm. The 60 μm sections were stained for NeuN (neuronal nuclei), which also labels neuronal cell bodies (Mullen et al., 1992), in order to determine the cortical layers and areal borders. The following protocol was used: Blocking and permeabilisation in 2 % PB-Tx (2 % TritonX-100 in 0.1 M phosphate buffer) with 10 % horse serum for 2 h at room temperature, followed by incubating the sections in the primary antibody mouse-α-NeuN (Millipore, MAB377, AB_2298772) diluted 1:500 at 4 °C for 2 days. The sections were then incubated with the secondary antibody donkey-α-mouse-AlexaFluor488 (Invitrogen, A-21202, AB_2535788, mice 1 and 3 to 5) or donkey-α-mouse-AMCA (aminomethyl-coumarin acetate, Jackson, 715-155-151, AB_2340807, mouse 2), diluted 1:500 for 6 h or overnight at 4 °C. Between antibody incubations, the sections were washed with 0.4 % PB-Tx, after the incubation with secondary antibody, the washes were done with PB only.

The BDA sections were washed in TB pH 8 (Tris buffer, 6.06 g Trizma.HCl and 1.36 g Trizma.Base in 1 l ddH2O). To visualize the BDA, the ABC kit (Avidin and Biotinylated horseradish peroxidase macromolecular Complex kit, Vector Lab) was applied. The sections were then incubated in 0.6 % (NH_4_)_2_Ni(SO_4_)_2_ * 6 H_2_O (ammonium Ni(II) sulfate hexahydrate) and 0.015 % DAB (3,3’-Diaminobenzidine) in TB pH 8 for 20 min. The catalyst H_2_O_2_ was added (to a final concentration of 0.005 %) and the reaction stopped after 3-5 min with the last washes in PB. The sections were mounted with DAPI (4’,6-diamidino-2-phenylindole, Vectashield mounting medium, H-1200, AB_2336790) onto slides for microscopy. (The AMCA-stained sections were mounted without DAPI because of the overlap in the emission frequencies of both fluorophores).

The BDA sections were mounted on gelatine-coated slides, air-dried and stained with Neutral red or Cresyl violet. For the staining, the sections were dehydrated and rehydrated in ascending and descending ethanol series. The tissue was incubated in Neutral red or Cresyl violet for ca. 7 min. Then followed a second dehydration series involving a differentiation step in 95 % ethanol acidified with acetic acid (ca. 1 %). After additional 15 min in 100 % ethanol, the sections were transferred to xylene and finally mounted with DPX. Overview micrographs of the sections were taken at 1.25x magnification and micrographs for counting and attribution to cortical layers and areas were taken at 10x magnification with a Leica DMRB fluorescence or light microscope and at 20x with a Zeiss Mirax Midi Slide Scanner.

### Analysis

The BDA-labeled brains were used for a qualitative comparison with the modified rabies virus label. The labeling pattern was very similar between the two tracers, but the BDA labeling was far sparser and only animals labeled with the more sensitive modified rabies tracer were analyzed quantitatively. The labeled cells were counted manually under the microscope and assigned to the corresponding areas based on the 10x and 20x micrographs. The identification of different cortical areas was based on detailed cytoarchitectonic characterization using the description of Caviness (1975) and the mouse brain atlas of Paxinos and Franklin (2001). The first step was to mark the boundaries of cortical layers based on cell density and soma size using the NeuN labeling. We used NeuN staining in addition to DAPI since it provides a clearer definition of layer and area borders, comparable to the Nissl staining. Boundaries were marked for every area of every section where a labeled cell was found, as well as in adjacent sections in order to confirm consistency of layer location. When cells were found exactly on layer boundaries, they were assigned to the lower layer in even sections and to the upper layer in odd sections. The boundaries between all cortical areas were then manually identified for each section in each animal by careful examination of the changes in cell densities and cortical layer thickness (as in Caviness, 1975).

Subcortical cells were mostly aligned to the mouse atlas, as the area boundaries could not always be determined cyto-architectonically. While functional or tracing methods offer the possibility of an even finer subdivision of cortical areas (e.g. Stiebler et al., 1997; Tsukano et al., 2015, for auditory cortex; and Wang and Burkhalter, 2007; Garrett et al., 2014, for visual cortex), they cannot offer the comprehensive assignment of every cell to a known location that can only be achieved with the classical cytoarchitectonic method.

Images taken with the Zeiss Mirax Midi Slide Scanner were exported with program PannoramicViewer. The different channels of the fluorescence micrographs were overlaid and brightness and contrast adjusted using GIMP (GNU image manipulation program, http://www.gimp.org, RRID: SCR_003182) or Adobe Photoshop CS6 (http://www.adobe.com/products/photoshop.html, RRID: SCR_014199). Using the Fiji plug-in TrakEM2 (Fiji, http://fiji.sc, RRID: SCR_002285, TrakEM2, http://www.ini.uzh.ch/∼acardona/trakem2.html, RRID: SCR_008954, Cardona et al., 2012), the micrographs of the labeled cells were manually aligned to the coronal sections of the mouse atlas and the volumes of the injection sites (i.e. the spread of the AAV) was measured. The number of cells per area and cortical layer was listed in a LibreOffice table, which was converted into a csv file. The data were then analyzed and graphed with Matlab (Mathworks, http://ch.mathworks.com/products/matlab.html, RRID: SCR_001622) or depicted with Adobe Illustrator CS6 (http://www.adobe.com/products/illustrator.html, RRID: SCR_014198) and Photoshop, respectively.

The fraction of labeled neurons (FLN, Markov et al., 2011) equals the fraction of labeled cells in a source area divided by the total number of labeled cells in the whole brain in percent. The extrinsic fraction of labeled neurons (FLNe, Markov et al., 2011) is the fraction of labeled cells in a source area divided by the total number of labeled cells excluding the injection site areas Au1 and AuV of the left hemisphere. For cortical areas, the fraction of labeled supragranular layer neurons (SLN, Barone et al., 2000) was calculated by dividing the number of neurons in layers 1-4 over the number of all labeled neurons in this area.

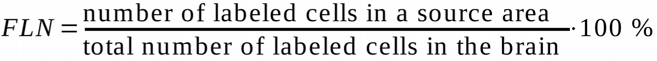

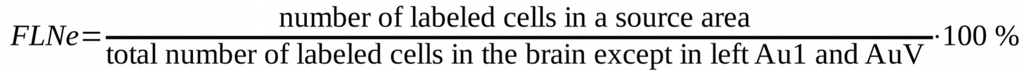

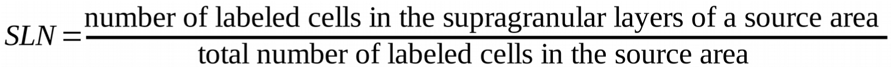

## Results

To obtain the complete input map of neurons connecting to Au1 from both hemispheres, single pressure injections of modified rabies virus together with AAV were made in Au1 of the left hemisphere in five mice. The location of the injection was confirmed in histological sections labeled with NeuN using cytoarchitectonical features to identify Au1. The injections sites were clearly visible under the epifluorescence microscope and confirmed that the modified rabies was injected through all layers of Au1 (see Figure 1A+B). The injection site volumes and locations were comparable between animals. The average injection site volume was 0.70 ± 0.19 mm^3^ and the virus reached a depth of approximately 0.9 mm (cortical depth of Au1 was 1 mm). At the center of the injection site, the distance from the rhinal fissure along the cortical surface to the center of the injection site was 1.44 ± 0.05 mm, which corresponded to a very consistent 20.3 ± 0.2 % of the total distance from rhinal fissure to the midline in each brain. Examples of coronal sections containing labeled neurons (in magenta) and stained for NeuN (in green) for cytoarchitectonic identification of cortical areas are shown in Figure 2A. Note the relatively dense labeling in the ipsi- and contralateral auditory cortices and in the medial geniculate nucleus of the thalamus, as well as distributed label in association cortical areas, and rhinal, visual, somatosensory and motor cortex.

**Figure 1.**
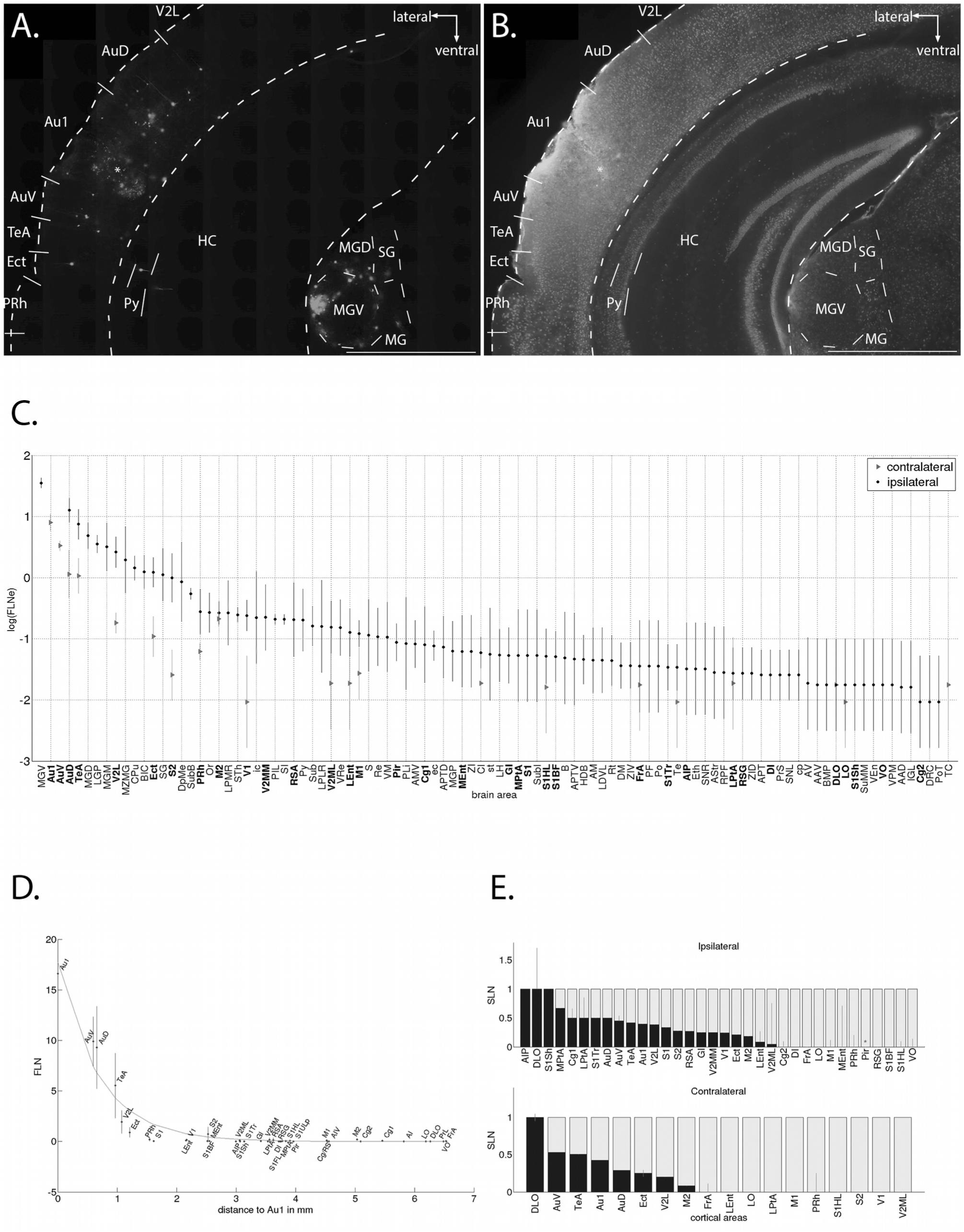
Example of an injection site. ***A*.** Coronal section through the injection site. The brightly labeled neurons have retrogradely transported the mCherry^+^ rabies virus to their somata and dendrites. Note the labeling in the cortex surrounding the injection site and in the thalamus. *B*. The same coronal section showing cells labeled with NeuN. At the injection site, the tissue is slightly damaged. The asterisk (*) denotes the center of the injection site. Scale bars 1 mm. ***C*. Mean FLNe (‘fraction of labeled neurons extrinsic’) values.** The FLNe, or input weight, is shown on a logarithmic scale for all brain areas with labeled cells. Cortical areas are written in bold. The projections from the contralateral hemisphere (grey triangles) are generally lower and sparser than from the ipsilateral hemisphere (black circles), but follow a similar pattern of relative weights. Error bars denote the relative standard deviation. ***D*. Distance-dependence of FLN of the ipsilateral cortex.** Mean FLN values sorted according to the distance of area centers to the injection site. Ca. 90 % of labeled neurons in the ipsilateral cortex were found within 1 mm distance, while the longest projections spanned 7 mm. Average FLN values and standard deviations are plotted as a function of distance to the primary auditory cortex. The curve is an exponential fit: FLN =17.76 * exp(-1.457 * distance) (R-square: 0.93). Areas above the curve show relatively stronger projection weights for their distance. Au1: primary auditory cortex. FLN: fraction of labeled neurons. ***E*. Mean SLN (fraction of labeled supragranular layer neurons) values of cortical areas.** Cortical areas were sorted based on the mean SLN values, from purely feedforward (SLN=1) to feedback (SLN=0), with standard deviations. Note that apart from three exceptions, all areas project in a lateral or feedback fashion to Au1, i.e. with SLN values between 0.5 and 0. Since the piriform cortex (Pir, marked by an asterisk) is part of the three-layered archicortex, its SLN could not be determined and was set to 0 because most of the labeled cells were found in the deep layer 3. Other areas without a dark bar had an SLN of 0. Areas without error bars were either represented by one animal only or had identical SLNs in the different animals. In the contralateral hemisphere, fewer areas are shown because fewer areas contained labeled cells. For area abbreviations see Table 1.

**Table 1:** Alphabetic list of brain area abbreviations based on the mouse brain atlas (Paxinos and Franklin 2001).

|  |  |
| --- | --- |
| PLi | posterior limitans thalamic nucleus |
| Po | posterior thalamic nuclear group |
| PoT | posterior thalamic nuclear group, triangular part |
| PRh | perirhinal cortex |
| PrL | prelimbic cortex |
| PrS | presubiculum |
| Py | pyramidal cell layer of the hippocampus |
| Re | reuniens thalamic nucleus |
| RPF | retroparafascicular nucleus |
| RSA | retrosplenial agranular cortex |
| RSG | retrosplenial granular cortex |
| Rt | reticular thalamic nucleus |
| S | subiculum |
| S1 | primary somatosensory cortex |
| S1BF | primary somatosensory cortex, barrel field |
| S1DZ | primary somatosensory cortex, dysgranular region |
| S1FL | primary somatosensory cortex, forelimb region |
| S1HL | primary somatosensory cortex, hindlimb region |
| S1Sh | primary somatosensory cortex, shoulder region |
| S1Tr | primary somatosensory cortex, trunk region |
| S2 | secondary somatosensory cortex |
| SG | supragenulate thalamic nucleus |
| SI | substantia innominata |
| SNL | substantia nigra, lateral part |
| SNR | substantia nigra, reticular part |
| st | stria terminalis |
| STh | subthalamic nucleus |
| Sub | submedius thalamic nucleus |
| SubB | subbrachial nucleus |
| SubI | subincertal nucleus |
| SuMM | supramammillary nucleus, medial part |
| TC | tuber cinereum area |
| Te | terete hypothalamic nucleus |
| TeA | temporal association area |
| V1 | primary visual cortex |
| V2L | secondary visual cortex, lateral area |
| V2ML | secondary visual cortex, mediolateral area |
| V2MM | secondary visual cortex, mediomedial area |
| VEn | ventral endopiriform nucleus |
| VM | ventromedial thalamic nucleus |
| VO | ventral orbital cortex |
| VPM | ventral posteromedial thalamic nucleus |
| VRe | ventral reuniens thalamic nucleus |
| ZI | zona incerta |
| ZID | zona incerta, dorsal part |
| ZIV | zona incerta, ventral part |

**Figure 2.**
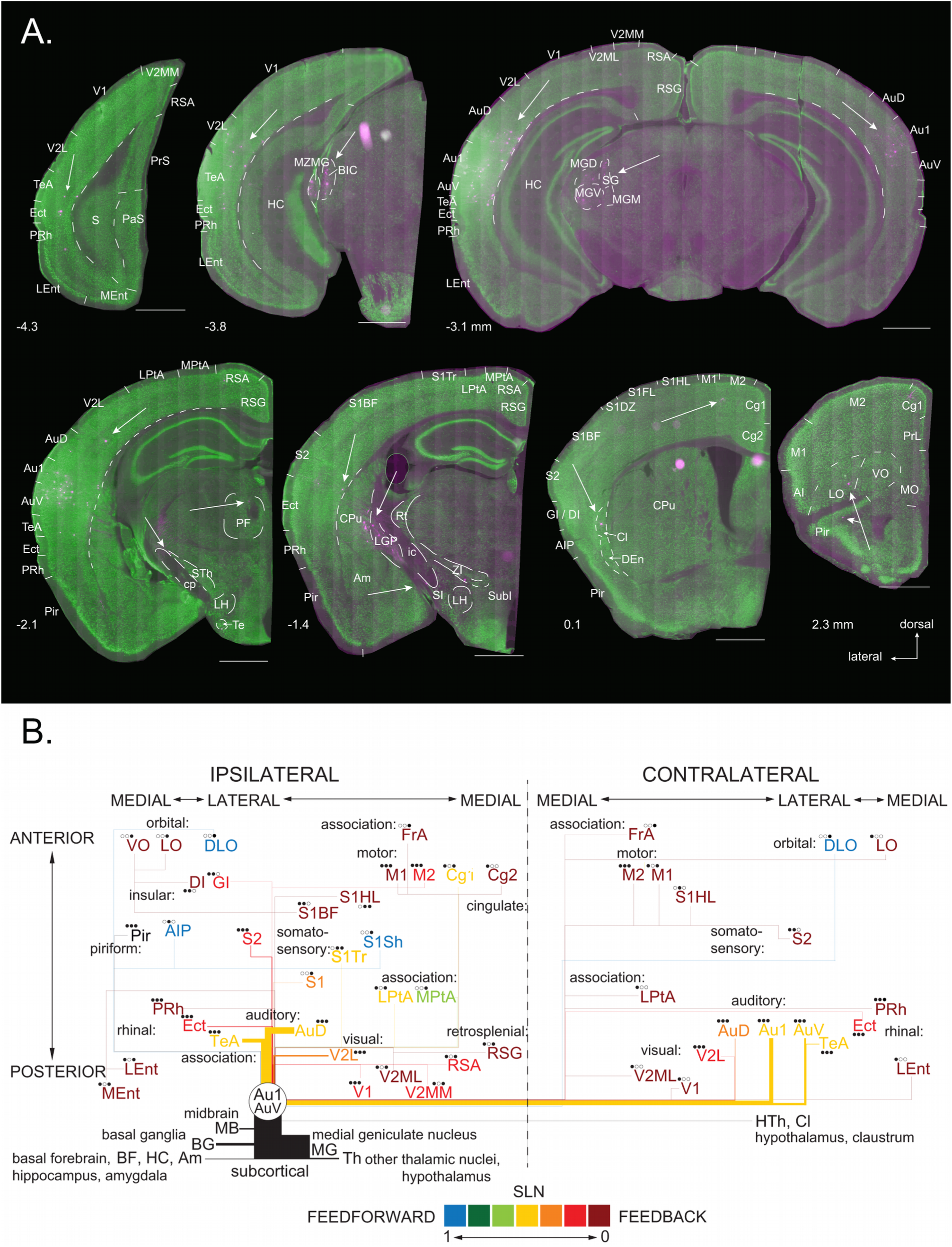
*A*. Distribution of labeled cells in one brain. Coronal sections containing labeled neurons (in magenta) and stained for NeuN (in green). Sections are ordered from posterior to anterior from the top left to the bottom right. Numbers below the sections correspond to the distance from Bregma (positive values are anterior from Bregma). If only one hemisphere is shown, it is the hemisphere ipsilateral to the injection. White arrows point out single cells or group of cells in temporal association area, rhinal, visual, auditory, somatosensory, motor and orbital cortices as well as in the inferior colliculus, the auditory thalamus and the striatum. Borders of subcortical areas are only delineated if they contain retrogradely labeled neurons or are adjacent to such an area. Am: amygdala; For other area abbreviations see Table 1. Scale bars 1 mm. ***B*. Overview of FLNe and SLN values.** The areas are arranged according to hemispheres and their approximate anterio-posterior and medio-lateral position. Colors indicate SLN (blue corresponding to feedforward and dark red to feedback connections), line thicknesses correspond to the FLNe. The dots represent different mice; filled dots indicate a labeled area in the corresponding mouse. FLNe: extrinsic fraction of labeled neurons. SLN: fraction of labeled supragranular layer neurons. For area abbreviations see Table 1.

Figure 1A+B shows an injection site of the viruses nine days after the injection. We coinjected AAV because after the long survival times we found that the rabies was far less visible at the injection site, which made the limits of the rabies injection difficult to demarcate. The extent of the rabies was confined to Au1, but the AAV filled most of Au1 and a portion of the secondary auditory cortex, AuV, which lies more ventrally. It was thus possible that some of the rabies particles had also reached the AuV and therefore we were conservative and assumed that the injection site included both Au1 and AuV. Thus when referring to projections to Au1, we implicitly include AuV. In one animal the tip of the pipette touched the hippocampus immediately below Au1, and the modified rabies labeled hippocampal neurons and possibly also a few of the projections to area CA1. This animal was excluded from the final quantitative analysis, although the pattern and weights of inputs to Au1 – after taking into account the intra-hippocampal connections – was similar to the brains where the injection was confined to the neocortex. In another animal, the injection labeled a small total number of neurons (ca. 1000), The major input sources of Au1 were the same as in the larger injections, but the numbers were too few to provide accurate counts and this was excluded from the quantitative analysis.

### Fraction of labeled neurons in extrinsic sources (FLNe)

The size of the pressure injection site was relatively large and covered more than half the volume of Au1, so although the fraction of labeled cells in Au1 was 26.5 ± 1.1 % of all labeled cells, this is likely to be an underestimate since we excluded cells within or at the border of the injection site that might have taken up the virus directly through their soma or dendrites. Thus to calculate the weights of the projections, we excluded the labeled neuron in Au1 and AuV and expressed the counts as a ‘fraction of labeled neurons extrinsic’ (FLNe), as a proportion of total labeled cells in the surrounding brain areas (Markov et al., 2011). The FLNe provides a simple measure of the numerical ‘weight’ of the many projections to Au1/AuV. The averaged FLNe values for the three mice were ranked by magnitude and plotted on a log axis (Figure 1C).

Thirty-four of the 47 neocortical areas identified contained neurons projecting to Au1. These areas were identified based on their cytoarchitectonic properties in NeuN-stained sections (see Methods for details). The most prominent labeling occurred in the auditory cortices (Figure 1C). The ipsilateral AuD (secondary auditory cortex, dorsal part) was the most densely labeled cortical area with an FLNe of 12.7 ± 5.7 % (i.e. mean ± standard deviation). The ipsilateral temporal association cortex (TeA, FLNe 7.5 ± 4.2 %) provided major projections to Au1/AuV. Although TeA is the association cortex that is physically closest to the auditory cortex, it was not the only association area to provide input to Au1/AuV, since the parietal association cortex (LPtA, MPtA, FLNe 0.08 ± 0.12 %) and even the distant frontal association cortex (FrA, FLNe 0.04 ± 0.06 %) projected to Au1.

A striking finding was the high degree of direct cross-modal projections to Au1, both in terms of projection weight and number of areas involved. The visual cortices were highly connected with Au1/AuV, having a cumulative FLNe of 3.3 ± 1.5 %. The lateral secondary visual cortex (V2L) contributed the biggest part, while the primary visual cortex (V1) and the medial secondary visual cortices (V2MM, V2ML) contributed less than 1 % each. Other sensory modalities formed weaker projections. The somatosensory cortex (primary and secondary) had a cumulative FLNe of 1.2 ± 1.0 %. Although the secondary somatosensory cortex (S2) provided a stronger projection than the primary somatosensory cortex (S1), labeled neurons were spread across the whole somatotopic representation in the primary somatosensory cortex (S1, S1BF, S1HL, S1Sh and S1Tr), with the barrel field (S1BF) being the strongest of these.

The olfactory system was also linked to the auditory cortex via a direct projection from the olfactory association cortex or piriform cortex (Pir, FLNe 0.09 ± 0.06 %), which is part of the three-layered archicortex. Weak projections also originated from the gustatory or insular cortex. Both granular and agranular parts of the insular cortex contained labeled neurons, but none of these regions were labeled in all animals, which is reflected in the low FLNe values (GI, DI and AIP, cumulative FLNe 0.11 ± 0.10 %).

The Au1/AuV not only received input from sensory cortical areas, but also from the primary and secondary motor cortex (M1, M2, FLNe 0.39 ± 0.08 %). These projections did not involve large numbers of neurons, but they were seen consistently in all brains. Moreover, the rhinal cortices of the parahippocampal region were the source of a substantial input to Au1. These regions included the ectorhinal cortex (Ect), lying on the ventral border of the temporal association cortex (TeA), and successively the areas of perirhinal (PRh) and lateral (LEnt) and medial (MEnt) entorhinal cortex. Altogether, the FLNe of the rhinal cortex was 1.7 ± 0.8 %. Sparser projections originated in the medial cortices: the cingulate and retrosplenial cortex had a cumulative FLNe of 0.32 ± 0.42 %. The labeling in the retrosplenial cortex (RSA and RSG, also called the posterior cingulate cortex), was denser than the labeling in the transition zone Cg/RS and in the cingulate cortex (or anterior cingulate cortex, Cg1 and Cg2). Also associated with the limbic system, the orbital cortex sent a weak connection to the auditory cortex. Lateral (LO), ventral (VO) and dorsolateral (DLO) orbital cortex contributed 0.05 ± 0.09 % to the total FLNe.

Most of the same cortical areas labeled in the ipsilateral cortex were also labeled in the contralateral cortex, but with a much smaller weight. The total FLNe of the contralateral cortex was 14.3 ± 3.4 %, whereas the FLNe of the ipsilateral cortex 27.4 ± 6.0 %. Note that the ipsilateral FLNe is an underestimate because the local intra-areal input around the injection site could not be included in this calculation. For single areas, the projection from the corresponding area in the contralateral cortex was in most cases much weaker. This difference in input weight between hemispheres was especially evident in areas with a relatively high ipsilateral FLNe value; the contralateral AuD, for instance, provided a tenfold weaker projection than the ipsilateral AuD. Eighteen cortical areas were labeled in the contralateral hemisphere and involved areas of the auditory, visual and somatosensory modality, motor and association cortex as well as rhinal cortex (Figure 1C; Figure 2B). Subcortical labeling, however, was very sparse in the contralateral hemisphere. Only three nuclei were labeled with 1 or 2 cells each, whereas on the ipsilateral side, labeled neurons were found in 63 different subcortical nuclei. These were mostly concentrated in the auditory thalamus, but spread as far as the cortical subplate and the deep midbrain. Overall, the FLNe from ipsilateral subcortical areas was 58.3 ± 3.3 %. Relatively strong projections were found from the basal ganglia, including the striatum (caudate/putamen, CPu, FLNe 1.44 ± 0.65 %), globus pallidus (LGP, MGP, FLNe 3.6 ± 1.1 %), substantia nigra and subthalamic nucleus. Of these areas, the lateral globus pallidus (LGP) had the most labeled cells, followed by CPu.

In the mice we injected with BDA, we also found widespread anterogradely labeled fibers in the caudate putamen and the lateral globus pallidus. In the mouse with a bigger BDA injection, there were also a few retrogradely labeled cell bodies inside these areas, but mostly close to the surrounding brain areas. Also some of the rabies-positive cells in the caudate putamen and the lateral globus pallidus were close the transition zones to the amygdala or the basal forebrain, but in addition many of the rabies-positive cells were clearly inside the striatum and pallidum. In contrast to the BDA-labeled axons, even the rabies-positive cells were restricted to the dorsal part of the pallidum and striatum, thus calling the connection between caudate putamen / globus pallidus and Au1 ‘reciprocal’ may not be strictly correct.

Other areas linked to the basal ganglia had a few labeled cells, namely the zona incerta (ZI) and the reticular thalamic nucleus (Rt). The zona incerta splits into a dorsal (ZID) and a ventral (ZIV) part, which are easily distinguishable as two parallel beams of cells in a coronal section. Together with the subincertal nucleus (SubI), the FLNe of the zona incerta was 0.18 ± 0.22 %.

Several nuclei of the basal forebrain projected to Au1, with a cumulative FLNe of 0.3 ± 0.1 %. They included the substantia innominata (SI), the basal nucleus (B) and the horizontal limb of the diagonal band (HDB). Of these areas, the substantia innominata was consistently labeled in all animals. Labeled areas in the cortical subplate involved the endopiriform cortex (VEn) and the claustrum (Cl), which projected to Au1/AuV from both hemispheres. With a cumulative FLNe of 0.08 ± 0.04 %, these projections were of a low weight.

Although the thalamus is expected to be one of the main input sources of a primary cortical area such as Au1, the projections from the thalamus were surprisingly numerous and together constituted close to half of the total extrinsic projection weights, i.e. 48.6 ± 3.8 % of the FLNe. The main contributor was the ventral medial geniculate nucleus (MGV – the auditory thalamus) with an FLNe of 35.5 ± 6.7 %. Surrounding this dense cluster of retrogradely labeled neurons, the other nuclei of the medial geniculate body (MGD, MGM, MZMG, SG) contributed 11.2 ± 4.4 % to the FLNe. The labeling in the thalamus extended beyond the medial geniculate body to various nuclei of the ventral, lateral and posterior thalamus. These thalamic nuclei typically had a projection weight of less than 0.3 %. In total, including the five areas of the medial geniculate body, 26 thalamic nuclei were labeled.

The midbrain also provided a number of projections. The most prominent of these was from the inferior colliculus (BIC) including the subjacent subbrachial nucleus (SubB), providing a total FLNe of 1.8 ± 0.7 %. A number of neurons projecting to Au1 were located in the deep mesenchymal nucleus (DpMe, FLNe 0.9 ± 1.3 %). The Au1 received direct input from nuclei known to modulate cortical activity such as the anterior pretectum and the dorsal raphe nucleus, but both areas provided a low input weight: The FLNe of the anterior pretectum (APTD, APTV, APT) amounted to only 0.15 ± 0.16 % and the dorsal raphe nucleus (DRC) had an FLNe of 0.01 ± 0.02 %. The substantia nigra, also a midbrain nucleus, sent projections from both the lateral (SNL) and the reticular (SNR) part with an FLNe of 0.06 ± 0.08 %.

A small number of neurons were consistently labeled in the hippocampus (Or, Py, FLNe 0.5 ± 0.4 %) and the subiculum (S, PrS, FLNe 0.14 ± 0.17 %). In the hippocampus the labeled neurons were mainly in the CA1 region close to Au1 in the oriens and pyramidal cell layers. These lay some distance from the injection site, indicating they were labeled retrogradely. The labeled hypothalamic cells were distributed among four nuclei, of which the lateral hypothalamic nucleus (LH) contained most cells. The cumulative FLNe of the labeled hypothalamic nuclei (LH, DM, Te, SuMM) was 0.14 ± 0.11 %. Finally, the amygdala complex was found to have a direct connection to Au1: from the anterior (AAD, AAV) and basomedial (BMP) amygdaloid nuclei as well as from the amygdalostriatal transition area (AStr). The FLNe of the amygdala was 0.08 ± 0.03 %, indicating a weak projection.

Across the whole brain, the FLNe spanned four orders of magnitude (Figure 1C). Areas with high FLNe values were consistently labeled in all mice. This was typically the case for FLNe values above 0.18 %, with two exceptions (DpMe and ic: internal capsule). Low FLNe values did not necessarily mean that the projections were variable, since several sources with an FLNe below 1 % were found in all mice, most notably the projection from the ipsilateral claustrum (FLNe 0.06 %). The strongest connection that was found only in one animal (from PLi, posterior limitans thalamic nucleus: FLNe 0.08 %) had a similar weight.

Of all the 98 areas with an observed projection to Au1, 30 were labeled in all animals and had a cumulative FLNe projection weight of approximately 96 %. Thirty-two areas were only found labeled in one animal. Most of these projections originated in subcortical areas and they typically contained only 1-2 cells per area. The cumulative FLNe of these uniquely labeled areas was less than 1 %.

Distance effects were clearly in evidence: the closer a cortical area was to Au1, the more labeled neurons it contained. For the ipsilateral cortex, the relationship of the FLN to the distance between the center of a projecting area and the injection site is plotted in Figure 1D. There was a correlation between the projection weights of the ipsilateral cortical areas and the distance of their centers to the center of Au1, but the rule is not absolute. Motor cortex (M2), for example, sends a stronger projection to Au1 than would be expected from its distance. Also the local input is well above the regression curve, while entorhinal cortices (LEnt, MEnt) and primary somatosensory cortices (S1, S1Sh, S1BF) show weaker connections than expected. Nonetheless, this distance effect is reflected in the fact that 89.0 ± 5.0 % of all labeled cells of the same cortical hemisphere were found within about 1 mm of the injection site. All these labeled cells lay in the four cortical areas closest to the injection site (Au1, AuV, AuD and TeA), whose centers were within 1 mm of the injection site. This shows the existence of a dense connectivity to areas around the auditory cortex and many sparse long-distance projections arising from all over the cortex in both hemispheres. The binary connection density of the ipsilateral cortex amounted to 72 % (34 out of 47 possible cortical areas were connected to Au1). The longest of these connections were about 7 mm, and originated from the ventral orbital (VO) and the frontal association (FrA) cortex (Figure 1D).

### Fraction of supragranular layer neurons in cortex (SLN)

The fraction of supragranular layer neurons (SLN) relative to all labeled neurons in a given area (see Methods) was calculated for all cortical areas. These values provide a means of ranking areas according to whether they were predominantly feedforward, lateral, or feedback relative to Au1 (see Barone et al., 2000). Here we found that Au1 received predominantly feedback connections (SLN < 0.33) from areas like visual, somatosensory, motor and rhinal cortex (see Figure 1E). This was also the case for primary somatosensory, insular and cingulate cortex, although the variability in the SLN of these areas was high due to the small numbers of labeled cells. Even in the piriform cortex, most labeled neurons were in the deepest layer, in layer 3, and it was thus classified as a ‘feedback’ area. Interestingly, SLN values in the midrange – between 0.33 and 0.66 – were common. Such values indicate ‘lateral’ projections, which arose from areas like the ipsilateral auditory cortex, the ipsilateral TeA and V2L, and from TeA, Au1 and AuV of the contralateral cortex (homotopic to the injection site). Other areas with an SLN around 0.5 (e.g. Cg1, LPtA, MPtA) were typically represented with few labeled cells and we interpret these SLN values with caution. The same caution applies to areas with an SLN of 1, where only a few labeled superficial layer neurons were found (AIP, S1Sh and DLO). With these exceptions, there were no projections with an SLN between 1 and 0.6, indicating that no cortical area was feedforward to Au1, as one might expect for a primary sensory area. On the other hand, the range of feedback and lateral projections (SLN between 0 and 0.5) was well-covered. Generally, the SLN did not differ substantially between the hemispheres, except that the proportion of labeled layer 5 cells tended to be higher in the contralateral hemisphere, at the expense of layer 6 cells. But this trend was obviously not reflected in the SLN. The data and analyses are given in a single comprehensive summary diagram (Figure 2B), which includes both the SLNs and the FLNs of the network that provides the many inputs of Au1/AuV.

## Discussion

Our analyses revealed that there are direct projections to Au1 from 98 brain areas of the mouse. The auditory cortex and the auditory thalamus provided the main weight in terms of numbers of projecting neurons, but the visual cortices, the temporal association cortex and the basal ganglia contributed substantially to the total input. Low-weight connections were found from almost every cortical area, the hippocampal formation, from many distributed nuclei in the basal forebrain, the hypothalamus, some nuclei of the thalamus, and the midbrain. Although the FLNe values varied between animals, the main inputs to Au1 were consistent. Large standard deviations were seen in areas with lower weights, simply because such areas did not contain labeled neurons in all animals. Inter-animal differences may be due to slight differences in the size and/or location of the injection site, or to actual connection weight differences between individuals.

### Comparison with other mouse connectome studies

The Allen Brain Institute has investigated mouse brain connectivity on a grand scale using recombinant AAV as an anterograde tracer (Oh et al., 2014). We searched this database (http://connectivity.brain-map.org/) for the brain areas where we found retrogradely labeled cells and checked if injection of AAV into these areas had resulted in anterograde axon labeling in Au1. Indeed almost all projections to Au1 were confirmed in wildtype C57BL/6J mice. Only a few areas that we found to project to Au1 were either not injected with AAV (e.g. FrA, MGP, SNL, fiber tracts) or could not be distinguished from nearby areas due to the injection size or precision (e.g. Cg/RS, MPtA, SubB, lateral thalamic nuclei, ZID, ZIV, TC, hippocampal layers, several amygdaloid nuclei).

A similar pattern of cortical sites projecting to Au1 is evident in the iConnectome map (http://www.mouseconnectome.org/), which is another growing database of mouse connectivity based on co-injections of anterograde and retrograde tracers (Zingg et al., 2014). Our findings agree with the high degree of connections between sensory modalities described in Zingg and colleagues’ (Zingg et al., 2014) medial sub-network of the mouse cortex. Additionally, we found direct projections from the lateral sub-network (piriform and insular cortex) to Au1, implying that these sub-networks do not operate independently.

The existence of direct projections from the caudate putamen and globus pallidus to the neocortex has been disputed (see e.g Divac et al., 1987), but consistent with our observations, the Allen brain atlas (http://connectivity.brain-map.org/) recorded that sparse anterograde labeling was indeed present in all layers of the ipsilateral Au1 and mostly in deep layers 5 and 6 in a mouse brain with AAV injection in the dorsal part of the caudate putamen. Interestingly, retrograde tracer studies in the cat point to the auditory cortex as one of the main targets of direct connections from striatum and pallidum to neocortex (Jayaraman, 1980; Oleshko and Maisky, 1993). Although some cells in the pallidum that project to the neocortex might be interpreted as belonging to a cell population of neighboring brain areas (Divac et al., 1987), this does not hold in all cases. In the rat for instance, a separate population of small globus pallidus neurons projecting to the frontal cortex was described that was distinct from the larger cholinergic cells along the boundaries of the globus pallidus (Van Der Kooy and Kolb, 1985). In the mouse, a similar picture emerges: After small tracer injections, only one-directional connections from the neocortex to the striatum were reported for the ultrasonic field of Au1 (Hofstetter and Ehret, 1992).

The iConnectome study was performed using conventional retrograde tracers (Cholera toxin B, Fluorogold), so the similarity in pattern of label suggests that the rabies virus is taken up by the same population of axons. The major advantage of the rabies viral tracers over other retrograde tracers is its sensitivity, but the negative aspect of virus tracing compared to conventional tracers is its cytotoxicity. We used the less cytotoxic AAV virus to give us an estimate of the likely maximum extent of the modified rabies uptake zone. Since the AAV reached the AuV ventral of the main injection site into Au1, we cannot completely exclude the possibility that some of the labeled cells also project to AuV. The dense labeling in the MGV, which mainly targets the primary auditory cortex (Winer et al., 1999), indicates that the rabies virus was largely confined to Au1.

### Comparing species

We found a lower proportion of labeled neurons (26.5 %) within Au1 than was reported for cat Au1 (60 %, Lee and Winer, 2011) or macaque visual cortex (80 %, Markov et al., 2011). This difference is likely due to the fact that the size of the injection site was large relative to the size of Au1 in the mouse. A different trend was observed for the main thalamic input, where we found 35.5 % of extrinsic projections coming from the MGV alone. The same projection in the cat was determined to be 18 % (Lee and Winer, 2011). Thalamic input to the macaque cortex appears to be even lower by comparison (1 % to V1, Markov et al., 2011). In the Mongolian gerbil, auditory subcortical afferents formed 27 % of the total (Budinger and Scheich, 2009), compare the 43 % we found in the mouse.

Overall, the range of FLNe values spanned four orders of magnitude in the mouse, while in the macaque it spanned five orders of magnitude (Markov et al., 2012). The FLNe of the ipsilateral cortex was exponentially dependent on physical distance: 90 % of these cells were within 1 mm to the injection site (compared to 75 % of labeled cells within ca. 1 mm and 95 % of labeled cells within ca. 2 mm of V1 in the macaque, (Markov et al., 2011)).

Small-world networks have been a popular graph-theoretical model of cortical networks (Watts and Strogatz, 1998; Bullmore and Sporns, 2009; Budd and Kisvarday, 2012). The typical features of such networks, viz. high local clustering and short path lengths due sparse long-range connections, optimize the integration and segregation of information and have been seen as suitable description for cortical networks. However, the high binary connectivity found in the long distance connections in macaque (66 %, Markov et al., 2012) and in the mouse (72 %) cortex is inconsistent with that of a classical small-world architecture, as emphasized by Markov and Kennedy (2013).

### Cortical hierarchy

The SLN provides an index of hierarchical distance (Markov and Kennedy, 2013). ‘Lateral’ connections, which indicate that the areas lie at the same level of the hierarchy (SLN ≈ 0.5), may thus have been expected from auditory cortices and even from ipsilateral V2L and V1, but surprisingly, the association area TeA (of both hemispheres) also connected to Au1 in a lateral fashion. As in the macaque, the immediately neighboring areas of Au1 had SLN values around 0.5. The SLN gradually declined for the more distant areas (Ect, V1, S2), but did not correlate with physical distance. Anterograde tracing with AAV virus from these areas providing lateral connections shows very sparse projections to Au1 (http://connectivity.brain-map.org). The axons innervate all layers, but are particularly sparse in layer 4. This projection pattern is evident even if only neurons in superficial layers or in layer 5 expressed the fluorescent label in a Cre-dependent manner (see http://connectivity.brain-map.org).

Long-distance feedforward paths from cortical areas to Au1 were notably absent, while feedback connections (from rhinal, insular, visual, somatosensory, motor and cingulate cortex) dominated. By contrast, after injections of a Cre+ pseudotyped lentivirus into the auditory cortex of transgenic (Cre-dependent tdTomato-expressing) mice, Nelson and colleagues found that retrogradely labeled cell bodies were located preferentially in superficial layers of M1 and distributed over superficial and deep layers of M2 (Nelson et al., 2013). In that same study, however, anterogradely labeled axons in Au1 terminated mainly in layers 1 and 6, which is a typical feedback pattern according to the criteria formulated by Felleman and Van Essen (Felleman and Van Essen, 1991). An attempt to reconcile these findings with our results may be the different sensitivity of the viral tracers used or the fact that Nelson and colleagues injected more dorsally in the auditory cortex. While cortical feedforward connections were mostly absent, the primary thalamic feedforward projection from the MGV had a very large FLNe (approximately 35 %). Together with the finding that thalamic axons target proximal dendrite segments (Richardson et al., 2009), our results suggest that in the rodent Au1 cortical processing is primarily feedforward-driven. This is in sharp contrast to the picture that has emerged from the cat and primate visual cortex, where thalamic input is a very small fraction (approximately 1 %) of all the excitatory input.

### Functional implications

Our results indicate that the auditory cortex is not the monosensory area its name implies, but that multimodal integration could occur even at this early stage of cortical processing. Multisensory integration is widespread in the neocortex (Schroeder et al., 2003; Clavagnier et al., 2004; Ghazanfar and Schroeder, 2006; Budinger and Scheich, 2009) and the connection between auditory and visual cortex has received particular attention. In humans, striking illusions like the McGurk effect demonstrate the interaction of auditory and visual information to form a consistent percept (McGurk and MacDonald, 1976). In the mouse, the projection from V2L was shown to modulate Au1 activity (Banks et al., 2011), but what role this visual feedback plays in auditory perception is still unclear. Activity in the monkey auditory cortex is suppressed before and during certain movements, like those involved in vocalization (e.g. Müller-Preuss and Ploog, 1981; Eliades and Wang, 2003). This motor-auditory crosstalk at the cortical level may be a crucial component of auditory processing even in the mouse, which has a rich vocal repertoire. The FLN values for many of these projections are very small, which raises the pertinent question of how these few fibers exert any significant effect in Au1. The possibilities are that they act in synchrony or form very specific connections with a subset of ‘pointer’ neurons that control the gain of the recurrent connections in the local microcircuit of Au1 (Douglas and Martin, 2007). The anatomical weight, as measured here in the FLN, has to be translated into a functional and dynamic projection weight, which of course may differ even between areas with the same FLNe value.

## Conclusion

Studies of anatomical ‘hardwiring’ provide valuable information about the network properties of interareal brain connections and provide a basis for interpreting the results of physiological and behavioral studies. The relative simplicity of measuring SLN and FLN provides a very powerful method of comparison between areas, systems, and species. Our quantification of input weights has revealed similarities to the macaque data in that the major cortical connections with auditory cortex come from adjacent areas. The pattern in the mouse also differs substantially from the macaque in having a much higher weight of connections from the thalamus and in receiving direct input from a larger proportion of cortical regions than is evident in the macaque. This difference may indicate constraints of processing in small brains, or adaptive specializations that are more species-specific.

## Other Acknowledgements

We are indebted to Botond Roska for advice and for providing the modified rabies virus. Imaging was performed with support of the Center for Microscopy and Image Analysis, University of Zurich.

## Role of Authors

All authors were involved in all aspects of the study.

